# Anatomical and functional studies of isolated vestibular neuroepithelia from Ménière’s Disease patients

**DOI:** 10.1101/2024.04.15.589685

**Authors:** Hannah R Drury, Melissa A Tadros, Robert J Callister, Alan M Brichta, Robert Eisenberg, Rebecca Lim

## Abstract

Surgical removal of vestibular end organs is a final treatment option for people with intractable Ménière’s Disease. We describe the use of surgically excised vestibular neuroepithelium from patients with Ménière’s Disease for 1) anatomical investigation of hair cell and nerve fibres markers using immunohistochemistry and 2) functional studies using electrophysiological recordings of voltage-activated currents. Our data shows considerable reduction in and disorganization of vestibular hair cells in the cristae ampullares while nerve fibres are in contact with remaining sensory receptors but appear thin in regions where hair cells are absent. Electrophysiological recordings of voltage-activated potassium currents from surviving hair cells demonstrate normal activity in both type I and type II vestibular hair cells. In addition, current-voltage plots from type I vestibular hair cells are consistent with the presence of a surrounding calyx afferent terminal. These data indicate surviving hair cells in Ménière’s Disease patients remain functional and capable of transmitting sensory information to the central nervous system. Determining functionality of vestibular receptors and nerves is critical for vestibular implant research to restore balance in people with Ménière’s Disease.

**Summary Statement:** This study shows, that while there is significant hair cell loss in Ménière’s Disease patients, surviving type I and type II vestibular hair cells have normal voltage-activated conductances.

## Introduction

Ménière’s Disease is an intractable condition of the inner ear, affecting both balance and hearing. Ménière’s disease most often affects one ear, although in 20% of Ménière’s Disease cases, it can be bilateral (Huang and Young, 2015). Diagnosis of Ménière’s Disease typically occurs sometime after the onset of symptoms, and is classified into two categories; definite and probable Ménière’s Disease by the International Classification of Vestibular Disorders. Definite Ménière’s Disease is diagnosed by an observation of episodic vertigo syndrome (20 minutes to 12 hours) with low to medium frequency sensorineural hearing loss and fluctuating aural symptoms (hearing, tinnitus, and/or fullness). Probable Ménière’s Disease is defined as episodic vestibular symptoms (vertigo or dizziness) and accompanied by fluctuating aural symptoms occurring in a period from 20 minutes to 24 hours (Lopez-Escamez et al., 2015). Ménière’s Disease onset typically occurs between 40 – 60 years of age with a slightly increased incidence in women (Kim and Cheon, 2020).

To date there is no cure for Ménière’s Disease. An excellent recent review describes in detail the various treatment options ranging from conservative to destructive that are currently used by five different international Ménière’s Disease guidelines (Mohseni-Dargah et al., 2023). In all instances, at the end stage of disease, the last remaining treatment option requires either the removal of the peripheral vestibular organs or resection of the vestibular portion of the vestibulocochlear nerve. Excision of the vestibular organs in patients with Ménière’s Disease is typically achieved by a translabyrinthine approach (Roberts et al., 2020). In surgical approaches involving the ablation of peripheral vestibular organs, drilling with irrigation and aspiration are characteristically used. However, for tissue used in this study, the superficial aspect of each canal was drilled and the membranous labyrinth and vestibular neuroepithelium are carefully removed via micro-hook or Fisch micro-raspatory. In previous studies, this tissue from Ménière’s Disease patients has predominantly been used for immunolabelling studies. However, due to the proximity of our research laboratory and physiology expertise, we were able to use Ménière’s Disease tissue samples for real-time electrophysiological experiments. To optimise the use of this valuable tissue, after electrophysiological recordings, samples were fixed and used for immunolabelling analysis. Here we describe the methods for using surgically removed Ménière’s Disease tissue for immunofluorescent labelling and electrophysiological recordings.

The majority of studies have used tissue samples following surgical removal, but some also used cadaveric tissue from patients with Ménière’s Disease (Stephenson et al., 2021, Calzada et al., 2012). Previous post-mortem examinations of vestibular neuroepithelium in Ménière’s Disease suggest hair cell loss is highly variable, with preferential loss of type II vestibular hair cells and Scarpa’s ganglion neurons (Tsuji et al., 2000). It is not known whether the surviving hair cells in diseased tissue are functional. A goal of this study was to show that surgically removed vestibular neuroepithelium can be used not only in anatomical but also functional studies. The majority of studies using adult human Ménière’s Disease vestibular tissue have been histological, characterising and quantifying neuroepithelial components including hair cells, supporting cells (Lopez et al., 2005), membranous labyrinth (Wackym et al., 1991), and basement membrane (McCall et al., 2009). Immunolabelling experiments describing proteins involved in endolymph recycling and extracellular matrix components, (Lopez et al., 2019, Asmar et al., 2018, Calzada et al., 2012) have also been undertaken.

More recently, studies have harvested inner ear tissue removed from patients with Ménière’s Disease or vestibular schwannoma for investigations of regenerative capacity (Taylor et al., 2015) and for gene transfer studies (Kesser et al., 2007). These studies focus on the use of gene therapies to replace lost hair cells due to aging, disease, and damage (Taylor et al., 2015). There are very few studies that have recorded functional activity from human vestibular hair cells. Approximately 25 years ago, there were two studies that successfully took whole-cell patch-clamp recordings from human vestibular hair cells, though these were from dissociated cells (Oghalai et al., 1998, Oghalai et al., 2000). In addition, the recordings were voltage-activated currents from hair cells of patients with vestibular schwannoma. More recently, this laboratory has electrophysiologically recorded from human fetal vestibular hair cells in a semi-intact neuroepithelial preparation which maintains the intracellular mileu (Lim et al., 2014). To date, there have not been any functional studies that have reported the viability of whole-cell patch-clamp recordings from vestibular hair cells from Ménière’s Disease samples. As described above, there is variability in survival of hair cells in Ménière’s Disease. However, we do not know if remaining hair cells remain functional, nor do we know whether vestibular hair cells that survive in Ménière’s Disease tissue are type I, type II, or both.

Overall, there were two aims of this study, 1) to investigate the distribution of hair cell specific marker, myosin VIIa and nerve fibre marker, neurofilament H, in the vestibular neuroepithelium from patients with Ménière’s Disease and compare this to normal human vestibular neuroepithelial tissue and 2) confirm the viability of recording functional responses of surviving hair cells from Ménière’s Disease neuroepithelium. Understanding whether surviving hair cells are functional is important to inform regenerative studies which target specific cell types for differentiation.

## Materials and Methods

### Ethical Considerations

All research in this study was approved by Hunter New England Health Ethics Committee and all experimental work was approved by University of Newcastle Institutional Biosafety Committee and followed the SRQR reporting guideline (O’Brien et al., 2014). We report on the feasibility of using diseased human vestibular tissue for electrophysiological recordings. Therefore, the results shown here are representative of the quality of data that can be acquired from tissue affected by Ménière’s Disease. Written consent was obtained from donors prior to surgery. The surgical and research teams worked independently. The surgical team was not involved in collection of electrophysiological data, and research team members were not involved in surgical procedures. Data shown are from Ménière’s Disease tissue that required surgical removal of vestibular organs for symptom treatment. Surgeries were performed in local Newcastle hospitals in New South Wales, Australia.

All patient data were de-identified and stored securely. Two-factor authentication was required by researchers to access patient data, which included, gender, date of birth, and length of time since diagnosis. No other personal patient data were collected.

### Tissue Collection and Transport

The translabyrinthine approach was used to remove vestibular organs (Roberts et al., 2020). Briefly, the superficial aspect of each semicircular canal was drilled to expose the membranous labyrinth. A Fisch micro-raspatory was used to retrieve the vestibular neuroepithelium from the ampullae of the semicircular canals and the utricle and saccule from the vestibule. The neuroepithelium was then collected in cold Liebovitz’s cell culture media (Life Technologies), which contained (in mM); 1.26 CaCl_2_, 0.99 MgCl_2_, 0.81 MgSO_4_, 5.33 KCl, 0.44 KH_2_PO_4_, 137.93 NaCl, 1.34 Na_2_PO_4_, pH 7.45, 305 mOsM). A research team member transported the tissue samples to the laboratory at The University of Newcastle. The samples were inspected for viability and used for electrophysiological and/or immunolabelling experiments, depending on the tissue quality. Fetal samples were collected and prepared as described previously (Lim et al., 2014).

### Tissue Preparation: Immunofluorescent labelling

Samples for immunofluorescent labelling were fixed in 4% paraformaldehyde (0.1M PBS) overnight and then washed in 0.1M PBS and stored in 0.1 PBS and 0.05% sodium azide until processed. Neuroepithelial tissue was cryoprotected in 30% sucrose (0.1M PBS) overnight and sections (20 – 50 μm) were cut by cryostat. Primary antibodies against hair cells (*myosin VIIa*; rabbit anti-myosin VIIa, 1:200, Proteus Bioscience), and afferent fibres (*neurofilament H;* chicken anti-neurofilament H, 1:500, Merck), were used for immunofluorescent labelling. Sections were incubated overnight in primary antibodies then washed (3 x 0.1M PBS) and incubated in secondary antibodies conjugated to Alexa-594 (1:200, Abcam) or Alexa-488 (1:200, Abcam) for 2 hours. DAPI (1:1000) was used to label cell nuclei. Sections were mounted using 0.1M PBS / 50% glycerol and cover-slipped. Images were captured using either a Nikon Eclipse EZ1 confocal microscope or Zeiss Axio Observer confocal microscope.

### Tissue Preparation: Electrophysiology

Tissue was prepared as previously described (Lim et al., 2014). In brief, bone debris and membranes overlying the vestibular neuroepithelia were removed using fine forceps and curved micro-scissors. This procedure was done in cold (4 °C) Liebovitz’s L15 cell culture media (Life Technologies, Australia). The individual vestibular neuroepithelia were further isolated, secured in place by a stringed harp in a recording chamber containing fresh, room temperature, L-15 solution. The recording chamber (volume = 1 ml, flow rate 2 ml / min) was perfused with oxygenated Liebovitz’s cell culture media.

### Electrophysiological Recordings

Borosilicate glass microelectrodes (3 – 5 MOhm; King Precision Glass Inc., CA, USA) were pulled using a Narishige electrode puller (PC-10) and filled with KGluconate internal solution (containing in mM; 42 KCl, 98 K.gluconate, 4 HEPES, 0.5 EGTA, 1 MgCl_2_, 5 Na.ATP, pH 7.3, 290 mOsM). Electrophysiological recordings were made by targeting cells under a Zeiss Axioskop 2 microscope using infra-red DIC optics. Recordings were made using an Axopatch 200B amplifier and data were acquired using Axograph X software (sampled at 20 kHz and filtered at 2 – 10 kHz). Whole-cell voltage-clamp protocols were used to characterise cell type. To determine cell type a type I hair cell protocol was first used to establish if the cell possesses the type I hair cells specific GK,L conductance (see inset Figure 3). If a cell did not have a GK,L conductance, a type II hair cell protocol was used (see inset Figure 4). This is identical to the type I hair cell protocol, with the exception that there is no hyperpolarizing step to -120mV (see inset Figure 3).

**Figure 1.**
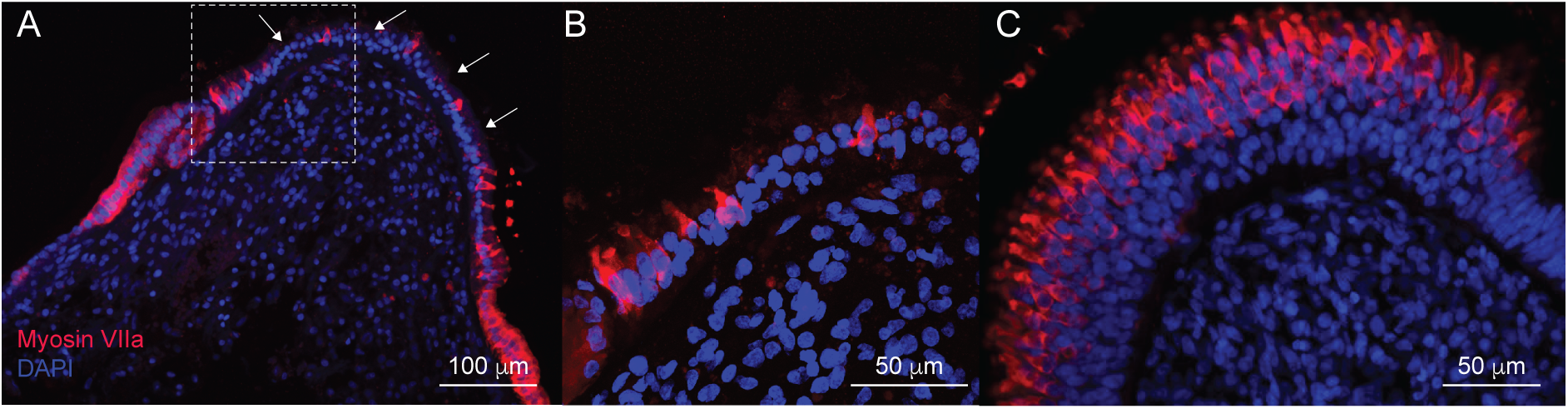
Loss of vestibular hair cells in posterior crista from Ménière’s Disease tissue. **A.** Immunolabelling with hair cell specific marker, myosin VIIa (red) shows few vestibular hair cells remain in the neuroepithelium. These are preferentially located at the periphery of the crista with few in the central zone of the crista. There are large voids where no hair cells are present (white arrows). Cell nuclei are labelled with DAPI (blue). **B.** High magnification image of the region within the dashed outline in A. Note, few hair cells are present and cell nuclei are disorganised within the neuroepithelium. A distinct supporting cell layer is absent. **C.** Myosin VIIa immunoreactivity in the crista of a developing human fetal inner ear (12 weeks gestation) shows a high density of vestibular hair cells (red) throughout the neuroepithelium.

**Figure 2.**
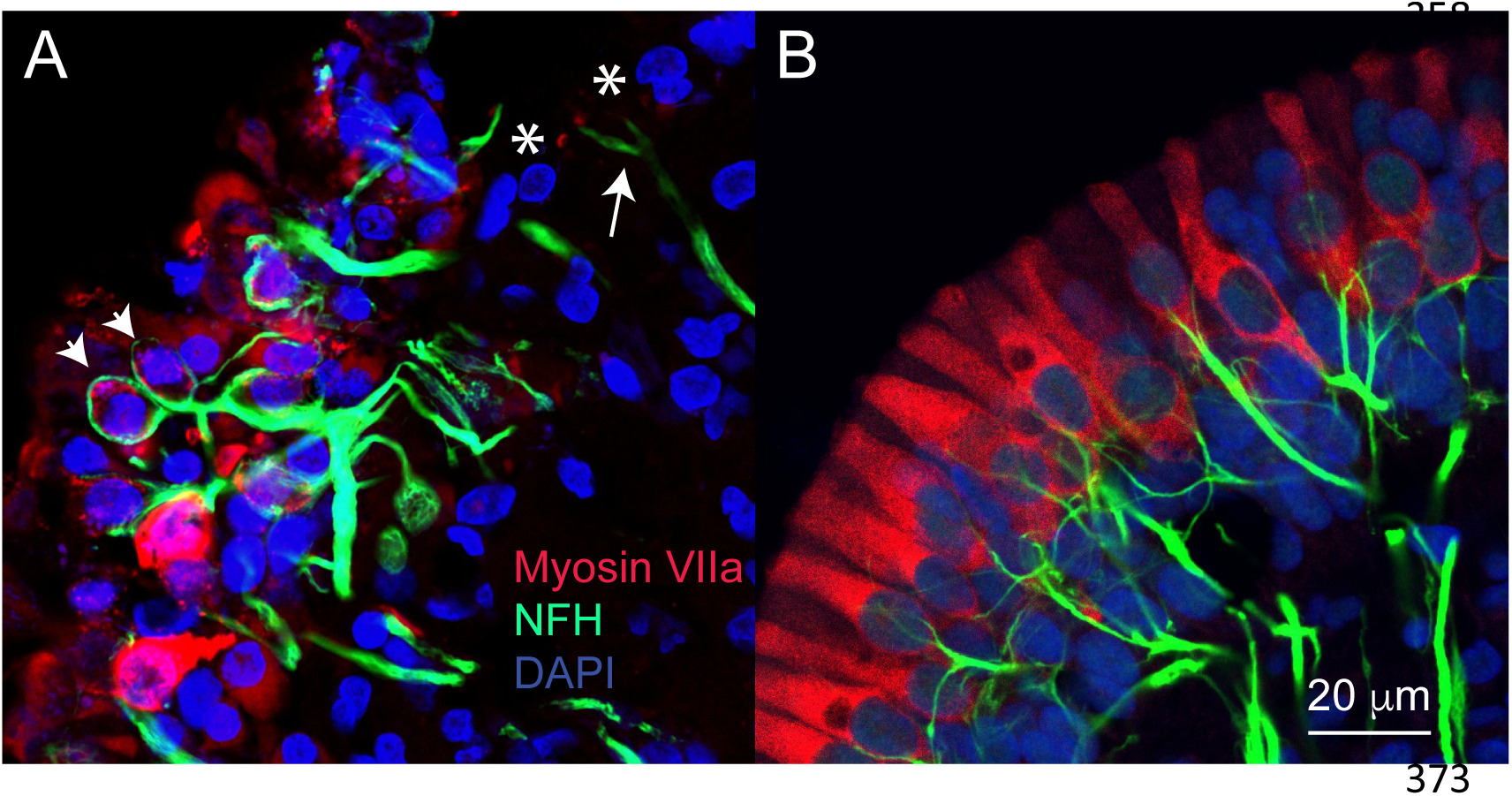
Immunolabelling of hair cells and nerve fibres in Ménière’s Disease neuroepithelium. **A.** Vestibular hair cells (red, Myosin VIIa) are disorganised within the vestibular neuroepithelium with some areas devoid of hair cells (white asterisks). Afferent nerve fibres (green, neurofilament H) are present within the neuroepithelium and in some instances appear to surround or encapsulate hair cells (white arrowhead). In regions of the neuroepithelium where hair cells are no longer present (asterisks), afferent fibres appear thinner (arrow). Cell nuclei are labelled using DAPI (blue). **B**. Hair cells (red) in the vestibular neuroepithelium of the fetal inner ear (14 WG) show an organised sensory monolayer with some distinctions in morphology beginning to emerge. In human fetal tissue, afferent nerve fibres (green) penetrate into the neuroepithelium and are beginning to contact hair cells. At this stage of development, no contacts yet resemble the characteristic type calyceal amphora morphology surrounding type I vestibular hair cells. Developing afferent fibres in the developing human neuroepithelium are thinner than those observed in the adult Ménière’s Disease neuroepithelium. Scale bar = 20 microns for panels A and B.

**Figure 3.**
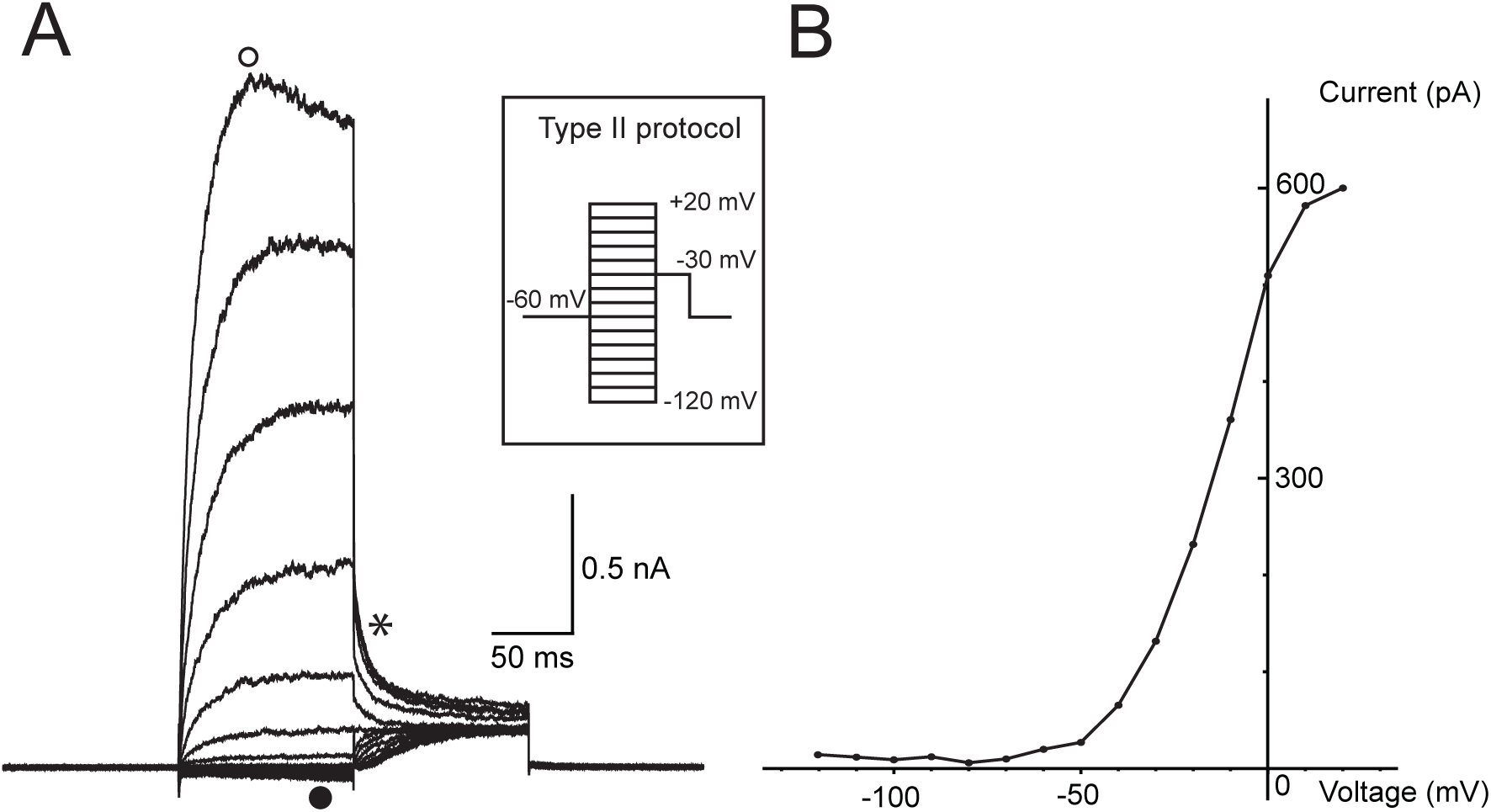
Whole-cell patch-clamp recording of a type II vestibular hair cell from Ménière’s Disease tissue. **A.** A voltage protocol (inset) is used to activate voltage-activated currents in a human type II vestibular hair cell from patient with Ménière’s Disease. This type II vestibular hair cell exhibited very small inward currents at hyperpolarized potentials (•) and large outward K^+^ currents at depolarized potentials (0). Instantaneous tail currents (*) were used to generate a I-V plot (B). **B.** The activation plot from tail currents of a type II vestibular hair cell shown in A. Fitting the I-V curve using the Boltzmann equation calculates the G_Max_ (maximum conductance) = 12.9 nS, V_½_ activation = -13.6 mV, and activation slope = 11.5.

**Figure 4.**
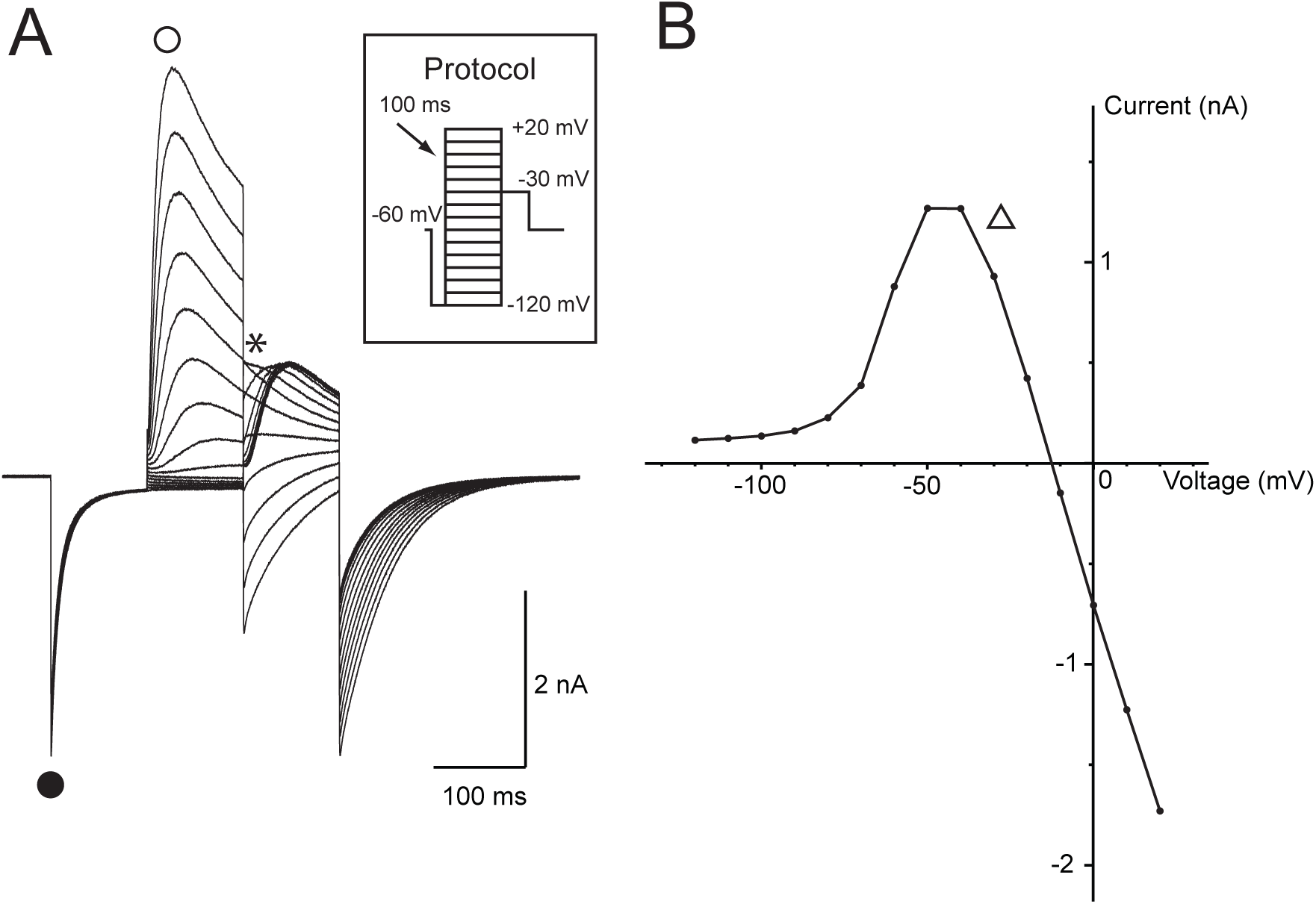
Whole-cell patch-clamp recording of a type I vestibular hair cell from Ménière’s Disease tissue. **A.** Inset shows voltage protocol used to activate voltage-activated currents in a human type I vestibular hair cell. This protocol delivers a -120 mV hyperpolarizing pre-pulse before the voltage ladder. This type I vestibular hair cell had large inward currents at hyperpolarized potentials below -70 mV (•), consistent with I_K,L_ and very large outward currents at depolarized potentials (0). Instantaneous tail currents (*) were used to generate an activation I-V plot (B). **B.** Activation I-V plot from tail currents of the type I vestibular hair cell shown in A. At voltages more positive than -40 mV, the currents begin to collapse (Δ) and reverse at ∼ -15 mV.

## Results

### Immunofluorescent labelling

Previous studies from surgically donated and cadaveric tissue have shown a loss of vestibular hair cells in patients with Ménière’s Disease (Ishiyama et al., 2007, McCall et al., 2009). Vestibular hair cell loss is highly variable between patients. Here we show immunolabelling of vestibular hair cells using myosin VIIa from a Ménière’s Disease patient. Figure 1 shows the posterior crista from a Ménière’s Disease patient. Figure 1A shows hair cells (red, Figure 1A and B) within the vestibular neuroepithelium. However, there are large voids, where there are no hair cells present (arrows, Figure 1A and B). This contrasts with normal developing human tissue where, even during early fetal development (12 weeks gestation), hair cells (red) are densely packed in the vestibular neuroepithelium (Figure 1C).

In Figure 2, immunolabelling of tissue from another sample with Ménière’s Disease shows disorganised hair cells throughout the neuroepithelium. There are regions where hair cells are absent (Figure 2A, asterisk) which contrasts with the dense packing of hair cells throughout the normal fetal neuroepithelium (Fig 2B). Ménière’s Disease tissue shows thick afferent nerve fibres (+) penetrating into the disorganised hair cell region terminating as endings that surround or enclose assumed type I hair cells (arrowheads). In contrast, in regions where hair cells are no longer present, the afferent terminals appear thinner (arrow; Figure 2A, asterisks). In the developing human fetal neuroepithelium, the afferent fibres appear thinner than the mature neuroepithelia of the Ménière’s Disease neuroepithelia. This is likely a function of development and at later stages of development, the fibres would increase in diameter. In the developing neuroepithelia, the afferent fibres are dispersed throughout the tissue and make contact with dense populations of hair cells (Fig 2B). At this stage of fetal development (14 weeks gestation), there do not appear to be calyceal terminals that surround type I vestibular hair cells.

### Electrophysiological recordings from type I and II vestibular hair cells

The immunolabelling results above (Figures 1 and 2), show there are significantly fewer vestibular hair cells in tissue from Ménière’s Disease patients compared to normally developing human neuroepithelia. This made targeting vestibular hair cells for electrophysiological recordings challenging.

### Type II vestibular hair cell

Whole-cell patch-clamp recordings were made from Ménière’s Disease neuroepithelium using the same technique as described in mouse (Lim et al., 2011) and developing human fetal vestibular neuroepithelium (Lim et al., 2014). Figure 3A shows a recording from type II vestibular hair cell. These hair cells do not possess the large low-voltage activated conductance (G_K, L_) that is characteristic of type I vestibular hair cells (see • Figure 4). Type II vestibular hair cells exhibit variable inward rectifying currents (•, Figure 3A) and large outward currents (0, Figure 3A) in response to the voltage protocol shown in the inset. These currents are similar to those previously observed in mice (Lim et al., 2011) and humans (Lim et al., 2014). The activation I-V plot (Figure 3B) for the cell’s instantaneous tail currents (*, Figure 3A) shows a characteristic sigmoidal I-V curve for a type II vestibular hair cell. Fitting the activation curve to this type II hair cells shows GMAX = 12.9 nS, V ½ = -13.6, and slope = 11.5.

### Type I vestibular hair cell

A type I vestibular hair cell in response to voltage protocol (inset, Figure 4A), shows large voltage activated currents at hyperpolarized potentials (•, Figure 4A), consistent with the presence of G_K,L_. During depolarizing steps, the type I vestibular hair cell also exhibits large outward currents (0, Figure 4A). These outward currents are significantly larger in amplitude than those recorded from the type II vestibular hair cell in Figure 3A. An activation plot (Figure 4B) taken from the instantaneous tail currents (*, Figure 4A) of the type I hair cell shows the presence of characteristic collapsing activation as the hair cell is depolarized above -50 mV. This differs significantly to the classic sigmoidal activation curve observed in type II hair cells and enzymatically isolated type I hair cells. We and others have shown previously that these ‘collapsing’ activation curves are a feature of type I hair cells that still remain surrounded by their associated calyx afferent terminal (Lim et al., 2011) (Contini et al., 2012). Potassium ions released from basolateral channels of the type I hair cells accumulate in the surrounding gap between hair cell and calyx terminal and significantly alter recording conditions. Thus, the presence of a collapsing activation curve in Figure 4 is indicative of a calyx afferent terminal surrounding a type I hair cell in the Ménière’s Disease sample. These collapsing tail currents are only observed in semi-intact neuroepithelial preparations where the intracellular milieu is maintained, and not in dissociated hair cell recordings.

## Discussion

In previous studies we describe the procedures for microdissection of vestibular neuroepithelium and removal of overlying membrane for its use in anatomical and functional experiments in mice and human fetal tissue (Lim et al., 2014, Lim et al., 2011). The same approach was used here. Our data show that using a translabyrinthine approach, the surgical removal of vestibular organs from Ménière’s Disease patients provides tissue that can be used for both immunolabelling and electrophysiological investigation. Indeed, given the surprising robustness and viability of Ménière’s Disease tissue for functional recordings, future work could also use cell culture techniques such as those previously described (Taylor et al., 2015) for additional longer-term investigations.

Consistent with previous anatomical data from deceased Ménière’s patients (Ishiyama et al., 2015, McCall et al., 2009), our samples of Ménière’s Disease vestibular neuroepithelia appear to exhibit the same severe hair cell loss and epithelia have become disorganised, which contrasts with the usual two distinct layers of 1) hair cells and 2) supporting cells.

Previous research showed the merging into a single disorganised neuroepithelial layer primarily in the cristae ampullares, but less so in the utricular maculae (Ishiyama et al., 2015). Immunolabelling studies using Ménière’s Disease tissue has shown distinct changes in expression levels of proteins involved in inner ear fluid homeostasis, proton channels, basement membranes, and extracellular matrix (Asmar et al., 2018, Calzada et al., 2012, Ishiyama et al., 2015, Lopez et al., 2019). Research suggests proteins such as aquaporins involved in inner ear fluid homeostasis are modified in Ménière’s Disease. Reports show altered expression of *aquaporins 4* and *6* in Ménière’s Disease neuroepithelium compared to acoustic neuroma and post-mortem normal tissue (Ishiyama et al., 2015). *Aquaporin 2* expression is elevated in the endolymphatic sac (Asmar et al., 2018), a site thought to be involved in development of endolympatic hydrops which is associated with Ménière’s Disease. *Cochlin*, a basement membrane component, was also significantly increased in Ménière’s Disease (Calzada et al., 2012). Together with our results, data shows there is significant remodelling of the vestibular neuroepithelium and altered expression of various proteins involved in maintaining fluid homeostasis and neuroepithelial structural integrity in Ménière’s Disease.

We show reduced labelling of afferent fibres by neurofilament H throughout the Ménière’s Disease mesenchyme and neuroepithelium. This is consistent with results described in a suspected Ménière’s Disease case which also show reduced afferent fibres (Okayasu et al., 2018). However, cross-sectional ultrastructural analysis of Ménière’s Disease nerve showed very low percentage of *abnormal* nerve fibres (Kitamura et al., 1997). Similar non-pathological results examining *ultrastructure* of vestibular nerve fibres observed at the level of the sensory epithelium and the internal acoustic meatus have been described (McCall et al., 2009). These ultrastructural studies however do not describe if there is an alteration in the number of afferent fibres in the neuroepithelium. Preliminary findings in this manuscript describe both reduced hair cell number <u>and</u> afferent fibres in Ménière’s Disease. It may be postulated that the absence of sensory input due to hair cell loss triggers the retraction and thinning of afferent fibres from the neuroepithelium. This would still be consistent with other studies that show varying degrees of degeneration and lack of abnormal or pathological nerve fibres that remain in contact with surviving hair cells (McCall et al., 2009).

Even with considerable hair cell loss and disorganisation of the vestibular neuroepithelium, some hair cells remain. These surviving hair cells appear resistant to the wholesale changes induced by Ménière’s Disease. We targeted these cells for electrophysiological characterization. Our data show hair cells from Ménière’s Disease tissue display characteristic voltage-activated currents consistent with type I and type II vestibular hair cells. GMAX, V_1/2_, and slope values for the type II vestibular hair cell are consistent with those previously recorded from human tissue, albeit foetal in origin (Lim et al., 2014). One other group have electrophysiologically recorded from adult human vestibular hair cells (Oghalai et al., 1998), although these were isolated vestibular hair cells, and none were from Ménière’s patients. We developed a semi-intact vestibular explant in mouse (Lim et al., 2011) and human (Lim et al., 2014) that maintains a more normal intracellular mileu, for electrophysiological recording of hair cells. Maintenance of this intracellular mileu, reveals a complex interaction of hair cells and their calyx afferent terminals (Lim et al., 2011, Contini et al., 2020). Our recordings from Ménière’s Disease tissue shows evidence of both type II vestibular hair cells and type I hair cell surrounded by a calyx afferent terminal. This suggests that Ménière’s Disease does not preferentially result in the death of a particular hair cell type. Gaining an understanding of whether the hair cell – afferent fibre connection remains functional in Ménière’s Disease is important for establishing which neural elements need to be targeted in regenerative and neuro-prosthetic studies. Recent regeneration studies have shown supporting cells in the vestibular neuroepithelium have the capacity for trans-differentation into functional type II vestibular hair cells, but there is no evidence for regeneration of type I vestibular hair cells (Hicks et al 2020, Gonzalez-Garrido et al.,2021). Further work needs to be undertaken to discover the signalling pathways that underlie the differentiation of vestibular type I hair cells that are essential for restoration of vestibular function. Most importantly however, we still need to determine the underlying cause of Ménière’s Disease. Without addressing the cause of Ménière’s Disease, treatment for this disorder including regeneration of vestibular hair cells, is likely to be compromised and will continue to induce severe vertigo and dizziness.

Further work remains to be done to establish the capacity of remaining vestibular afferent fibres to fire action potentials in the presence of Ménière’s Disease. If vestibular afferent fibres remain viable, there is enormous potential to stimulate these fibres using a vestibular implant in the same manner as cochlear implants stimulate cochlear afferent fibres to restore hearing. The ability to restore balance function for people with Ménière’s Disease would result in a significant improvement in quality of life for people with this devastating disorder.

## Acknowledgements

The authors would like to thank the donors for their valuable contribution to this work.

